# Stretchable, nano-crumpled MXene multilayers impart long-term antibacterial surface properties

**DOI:** 10.1101/2023.01.23.525034

**Authors:** Neha Nagpal, Mohammad Asadi Tokmedash, Po-Yen Chen, J. Scott VanEpps, Jouha Min

**Author notes:** These authors contributed equally to the manuscript (Neha Nagpal, Mohammad Asadi Tokmedash). Corresponding author: Jouha Min, PhD, Department of Chemical Engineering, University of Michigan, 2800 Plymouth Rd., Ann Arbor, MI, 48109.

## Abstract

Infections are a significant risk to patients who receive medical implants, and can often lead to implant failure, tissue necrosis, and even amputation. So far, although various surface modification approaches have been proposed for prevention and treatment of microbial biofilms on indwelling medical devices, most are too expensive/complicated to fabricate, unscalable, or limited in durability for clinical use. Here we present a new bottom-up design for fabricating scalable and durable nano-pattered coatings with dynamic topography for long-term antibacterial effects. We show that MXene layer-by-layer (LbL) self-assembled coatings -- with finely tunable crumple structures with nanometer resolution and excellent mechanical durability -- can be successfully fabricated on stretchable poly(dimethylsiloxane) (PDMS). The crumpled MXene coating with sharp-edged peaks shows potent antibacterial effects against *Staphylococcus aureus* and *Escherichia coli*. In addition, we find that on-demand dynamic deformation of the crumpled coating can remove ≥99% of adhered bacterial cells for both species, resulting in a clean surface with restored functionality. This approach offers improved practicality, scalability, and antibacterial durability over previous methods, and its flexibility may lend itself to many types of biomaterials and implantable devices.

## INTRODUCTION

Microbes have remarkable capabilities to adhere to abiotic surfaces and form multi-cellular communities called biofilms, which can negatively impact the safe use and function of medical devices in humans. Despite the advances in material science, implant-related infections remain one of the most devastating complications^[1]^. Release-based antimicrobial technologies using antibiotics^[2]^ or nanoscale antimicrobial substances (*e.g*., Ag, ZnO, Cu, Au) have been developed extensively^[3]^, yet there remain limitations to all commonly used antimicrobial surface modifications, such as loss of efficacy over time (due to depletion of eluting therapeutic agents) or systemic toxic effects on the patient. Furthermore, the development of antibacterial resistance and the emergence of superbugs, such as methicillin-resistant *Staphylococcus aureus* (MRSA) and extended-spectrum beta-lactamases (ESBL) producing bacteria, have resulted in new difficulties in the prevention and treatment of implant-related infections. With the growing use of biomaterials and implantable devices, there is a clear need for the development of multifunctional surfaces with long-term antibacterial activity.

A promising alternative to the conventional release-based approaches is surface topographical modification of the material, which can impart bactericidal or antifouling capabilities to prevent bacterial cell attachment or offer mechanical killing properties to the substratum. Previous research demonstrated that plants and animals had developed antibacterial micro- and nano-topographies that can be mimicked for antibacterial activities^[4]^. For example, nano-scale topographies such as those on the wings of cicadas and dragonflies have antimicrobial activities and can be mimicked for bacteria killing (*i.e*., bactericidal) and fouling control^[5]^. In general, nano-scale topographies have been shown to exhibit mechano-bactericidal capability by directly damaging the cell membrane^[4c, 5–6]^. On the contrary, micron-scale topographies do not typically have bactericidal effects but may inhibit bacterial adhesion (*i.e*., antifouling) through specific effects on cell-material interactions^[7]^. Because these topography-based physical approaches obviate the need for antimicrobial agents, the surface modification strategy will not promote the development of antimicrobial resistance associated with chemical approaches (*e.g*., antibiotic eluting surfaces). In recent years, many studies have been conducted to investigate the effects of nano- and micro-structured surfaces, with either well-defined or random size and distribution of features, against different species of bacteria^[8]^. Most studies to date, however, have been focused on static topographies, which have limitations in long-term biofilm control because even a small number of cells that attach can grow and gradually overcome the unfavorable topographies^[7a]^. Furthermore, the current micro-/nano-patterning techniques such as lithographic patterning of nanogratings are complicated, expensive, and not readily scalable, which hamper the translation of the research findings into practice.

Motivated by these challenges, we sought to develop a new strategy (**Scheme 1**) for long-term antibacterial activity through the integration of nano-pattered surfaces with dynamic topography. Specifically, for surface patterning, we used an emerging method that creates 3D crumple structures by controlled shrinkage of a stiff coating on a soft, compliant substrate^[9]^. Because of the modulus mismatch between the coating and the substrate, the 1D or 2D shrinkage result in wrinkled or crumpled structure^[10]^. Compared to the conventional lithographic methods, this method has advantages in the simplicity and scalability of fabrication. An exciting new approach to the creation of these patterned surfaces is the deposition of 2D sheet-like nanomaterials, such as MXene—a family of transition metal carbides/nitrides/carbonitrides—whose atomically thin nature enables the creation of ultrathin flexible films suitable for finely controlled patterning. Topographically patterned MXene has found numerous applications in optical and electronic devices^[11]^, soft robotics^[12]^, energy storage^[13]^, and functional coatings^[10]^. The usage of MXene in the biomedical fields is blooming recently, mostly in cancer therapies^[14]^, bone regeneration^[15]^, and antibacterial treatments^[16]^, but MXene as a biofilm interfering agent or nano-/micro-patterned MXene coatings as antibacterial surfaces have rarely been reported. For the coating deposition, we used the enabling nanofabrication tool of electrostatic layer-by-layer (LbL) assembly to create conformal nanoscale MXene multilayer coatings. The LbL method offers myriad advantages, including nanoscale control (down to a sub-nm resolution), conformal and surface-agonistic nature, scalability, and simple and low-cost processing^[17]^, which make it a most appropriate fabrication method for our purposes. For dynamic topography, we use PDMS substrates because of its stretchability (elasticity) and biocompatibility. PDMS has been widely used as wearable and implantable materials^[18]^. Based on these premises, we hypothesize that (1) nano-crumpled surface topographies, mimicking those on the cicada wings, can prevent bacterial attachment and (2) physically kill bacterial cells that encounter the surface; and (3) dynamic deformation in response to external stimuli (*e.g*., mechanical stretching, magnetic actuation) can release the debris of dead bacterial cells to (4) preserve this multifunctionality of the surface over the long-term.

**Scheme 1.**
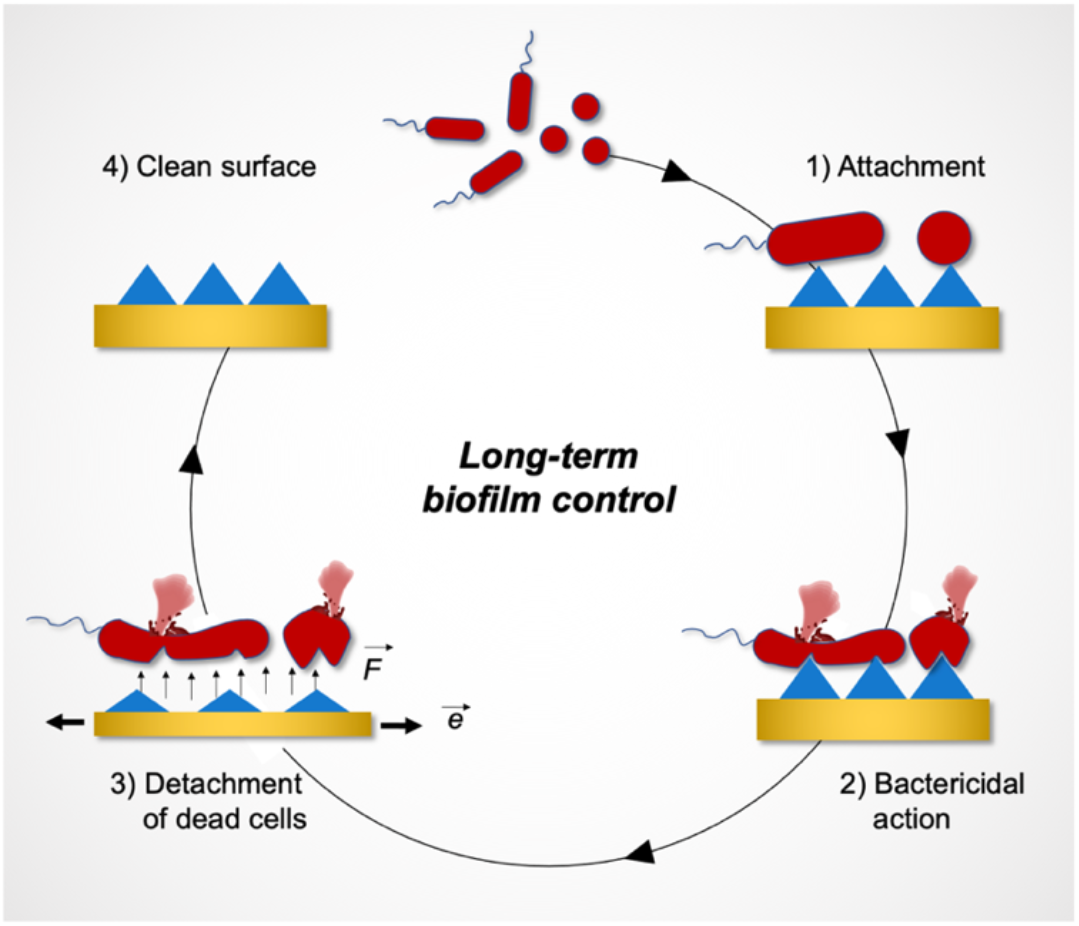
Scheme 1. Proposed mechanism of the stretchable, nano-crumpled (MXene/GS)_*n*_ multilayers for long-term biofilm control. **1**. Following *in vivo* implantation, opportunistic bacteria at the surgical or wound site can readily attach to biomaterial surfaces and cause infection. In an uncoated planar surface, the attached bacteria can readily form biofilm. **2**. In our LbL system, however, the nano-structured surfaces composed of sharp nano-edges of MXene nanosheets can induce bacterial cell death by damaging the bacterial membrane. **3**. The subsequent application of external stimuli (*e.g*., mechanical stretching to a prescribed strain, 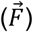) can induce dynamic surface deformation, which exerts enough force (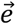) to release the debris of dead bacterial cells, **4**. resulting in a clean surface with restored multifunctionality.

In this study, we demonstrate a bottom-up design for fabricating stretchable Mxene multilayers with tunable crumple topography and explore the interaction of the finely tuned patterned surfaces with different species of bacteria (*i.e., S. aureus* and *E. coli*). To our knowledge, this will be the first demonstration of (i) implementing LbL assembly method to fabricate and control the crumpled structures of MXene multilayer coatings and (ii) the antibacterial and antibiofilm activity of highly crumpled, stretchable MXene multilayer coatings. Briefly, we first generated a conformal MXene multilayer coating via the LbL assembly method on a biaxially-oriented polystyrene (PS) substrate, and the as-prepared MXene coating was shrunk by thermal annealing to form a crumpled structure due to the shrinkage of the PS substrate. Subsequently, the crumpled MXene coating was transferred onto a flexible PDMS substrate by spin-coating PDMS directly on the crumpled MXene coating followed by the removal of the PS substrate. The morphological features of the crumpled MXene coating were characterized using scanning electron microscopy (SEM) and atomic force microscopy (AFM). The antibacterial effects of the coatings were evaluated by fluorescence microscopy and SEM. We further implemented surface deformation by mechanically stretching the MXene coatings to detach the debris of dead bacterial cells, and then examined the antifouling effects by fluorescence microscopy. Our data demonstrate successful biofilm treatment and removal through a combination of mechano-bactericidal and antibiofouling properties, without the release of any pharmacologic agents. The nano-/micro-structure of the surfaces and the dynamic deformation (*e.g*., strain, frequency) are tunable, and the fabrication process is scalable for different materials and substrates.

## RESULTS

### LbL assembly of (MXene/GS)_*n*_ multilayers

We fabricated the (MXene/GS)_*n*_ multilayer films (**Fig. 1a**) by sequential LbL deposition of positively-charged gentamicin sulfate (GS), and aqueous dispersions of negatively-charged Ti_3_C_2_T_x_ MXene nanosheets, produced by selective etching of aluminum atoms from Ti_3_Al_2_C_2_ MAX phase in a solution of HCl and LiF mixture (see Methods and **Fig. S1**)^[19]^. The multilayer films are denoted (MXene/GS)_*n*_ where *n* corresponds to the number of bilayers that are repeated in the LbL deposition process. We selected Ti_3_C_2_T_x_ MXene nanosheets as the building block units for LbL fabrication and subsequent nanomanufacturing due to their high-aspect ratio with atomic thickness, high charge density, high water solubility, proven antibacterial properties and high mechanical properties, which make it a most appropriate material for our purposes. MXene nanosheets in a stable aqueous dispersion at pH 6.1 had a highly negative charge (−37 mV by zeta potential). Transmission electronic microscope (TEM) images showed a lateral size of MXene sheets in the range of several hundreds of nanometers (**Fig. S1b**). We selected GS—a common aminoglycoside antibiotic—as the complementary species of MXene in the LbL assembly for several reasons: (i) highly positive charge due to the protonation of multiple primary amino groups at the fabrication condition (pH 5), (ii) small size (477 Da), (iii) broad-spectrum bactericidal activity against many Gram-negative and Gram-positive species^[20]^, and (iv) low cytotoxicity (no apparent cytotoxicity associated with GS up to 500 μg/mL, (**Fig. S2**). The high charge density promotes self-assembly processes (**Fig. S3**), especially, LbL self-assembly as it provides stronger electrostatic interactions between GS and MXene. The protonated amine groups of the GS would have a high affinity with the terminal group-doped metallic surfaces of MXene nanosheets and may even form chemical bonds^[21]^. We tested other antibiotics (*e.g*., vancomycin) as the counter phase for LbL assembly of MXene multilayers and found that agents with only one or two protonated amine groups do not self-assemble with MXene nanosheets.

**Figure 1.**
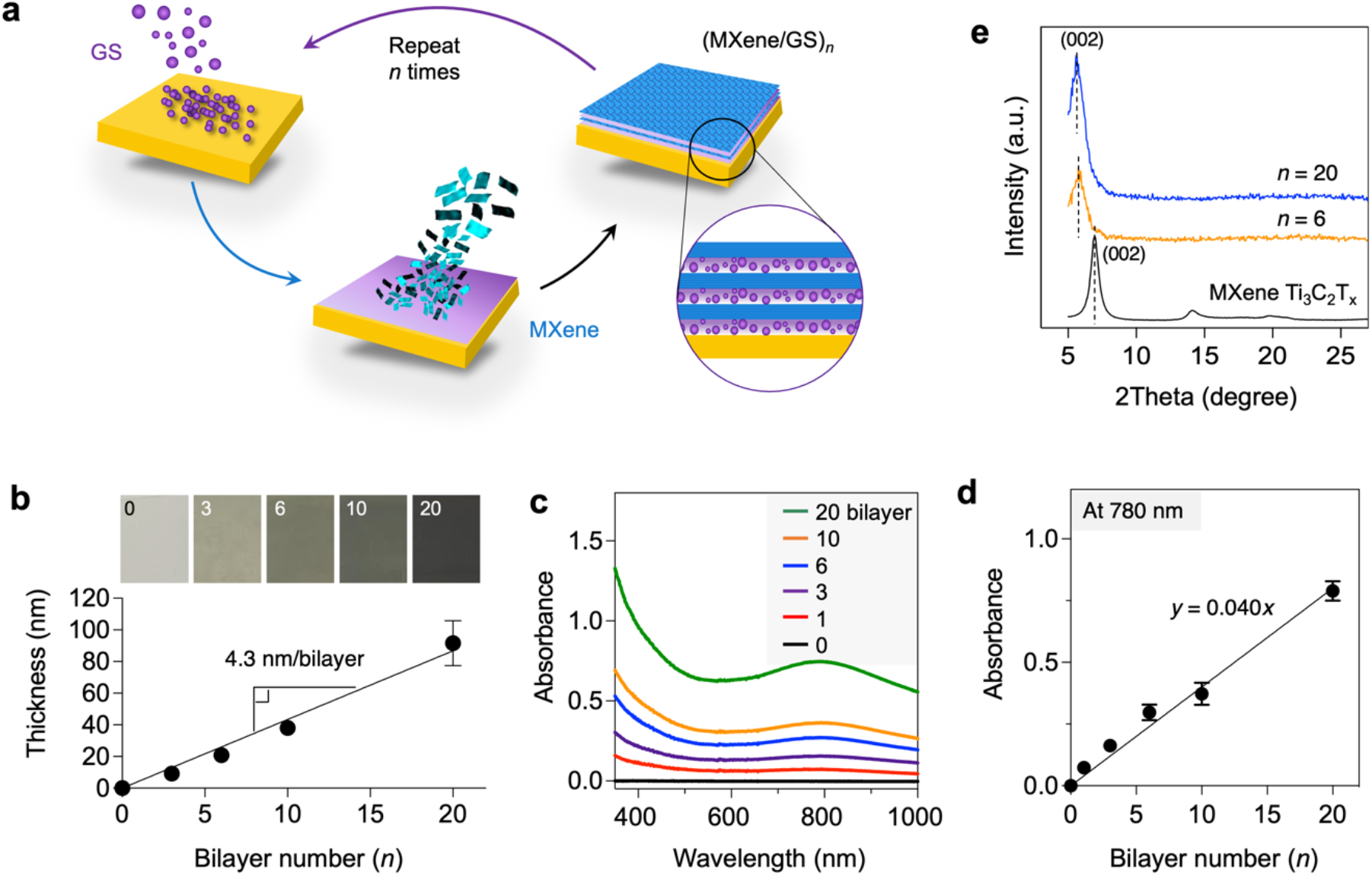
Structural characterization of as-prepared (MXene/GS)_*n*_ multilayers. **a**. Schematic of LbL nanofabrication of (MXene/GS)_*n*_ multilayers on biaxially-oriented PS films. **b**. Digital photographs of (MXene/GS)_*n*_ on PS films and growth curve of the films as a function of bilayer number (*n*) measured by AFM. **c**. UV-vis absorption spectra of (MXene/GS)_*n*_ on glass. **d**. Absorption values of (MXene/GS)_*n*_ vs *n* at 780 nm. **e**. XRD patterns of (MXene/GS)_*n*_ vs. *n*, and pure MXene film.

To investigate the growth behavior and scalability of LbL MXene multilayer films, we used an immersive-LbL self-assembly technique to grow the (MXene/GS)_*n*_ on large-area planar polystyrene (PS) sheets, glass, and silicon wafers. The color of the coating, which arises from the MXene sheets, became successively darker as the number of bilayer cycles increased from 1 to 20 (**Fig. 1b**). We measured the thickness of (MXene/GS)_*n*_ multilayers from the AFM images of scratched films grown on silicon wafers (**Fig. 1b**), which showed a linear increase in thickness with the bilayer number “*n*”. For a 20-layer-pair coating made by immersion, the thickness was 91.6 ± 14.2 nm. The coating’s UV-vis spectra demonstrate broad adsorption at 780 nm, consistent with the MXene nanosheets. The adsorption at 780 nm also increased linearly with *n* (**Fig. 1c-d**). The linear growth is a feature of a successful LbL self-assembly, and indicates that the two species completely alternate during the LbL process.

XRD patterns of the (MXene/GS)_*n*_ multilayers (**Fig. 1e**) showed ordered structures similar to pristine MXene films^[22]^, and the intensity of (002) peaks, stemmed from the ordered stacking of MXene sheet basal planes, increased with the number of bilayers, *n*. It should be noted that an ordered LbL structure was obtained only when the counter phase was the small-molecule GS, compared with reported polymer-based LbL structures such as (MXene/PDAC)_*n*_ and (MXene/PEI)_*n*_ films whose (002) peak completely disappeared due to their less ordered structure^[23]^. We attribute the ordered structure to the small size of GS which forms a sub-nanometer gap in between MXene sheets in the LbL films, leading to a quasi-intimate interfacial contact between the nanosheets similar to pure MXene films. Additionally, (002) peaks of (MXene/GS)_*n*_ shifted from 6.9° for pure MXene films to 5.8°, which showed a uniform pillaring effect of GS in LbL process. This corresponds to an increase of 2.4 Å in the average interlayer spacing from 12.8 Å up to 15.2 Å, which means an average interlayer distance of 1.52 nm between the MXene sheets in the multilayer films. Using the LbL approach, we could control the thickness of MXene multilayers at a resolution of few nanometers, which is essential for fine tuning of the feature sizes (wavelength and amplitude) of the crumpled MXene multilayers.

### Tunable crumpling of the MXene multilayer coating

**Figure 2a** illustrates the fabrication process of the stretchable (MXene/GS)_*n*_ multilayers with crumpled structure. After fabricating a MXene multilayer film on a PS substrate, we introduced a quick thermal annealing at 130°C, which is higher than the glass transition temperature of PS (~100°C), to shrink the substrate by releasing the prestretched strain. Because of the modulus mismatch between the MXene multilayer coating and the PS substrate, the 2D deformation resulted in isotropically crumpled MXene LbL films. The strain is defined as ε = (*L*_i_ – *L*_f_)/*L*_i_, where *L*_i_ are *L*_f_ are the lengths of the PS films before and after the shrinkage process, respectively. These wrinkles can further be transformed into ridge-like structures at high shrinkage rates^[24]^. Subsequently, we spin-coated PDMS on the crumpled MXene LbL film, and then dissolved the PS substrate in dimethylformamide (DMF) to create a stretchable MXene LbL film with the same crumpled structures^[11b, 25]^. Although MXene LbL multilayers can be directly incorporated onto a PDMS substrate, we implemented the two-step fabrication process (*i.e*., LbL assembly followed by transfer printing) because it was not feasible to create nanoscale sharp crumpled structures using the MXene multilayers on a prestretched PDMS, even at 1 bilayer (thickness ~4 nm). Compared to PS, PDMS can undergo only up to ~30% biaxial deformation strain and has the lower elastic modulus (0.5-1.5 MPa for PDMS vs. ~3.4 GPa for PS), which makes it difficult to create highly crumpled structures.

**Figure 2.**
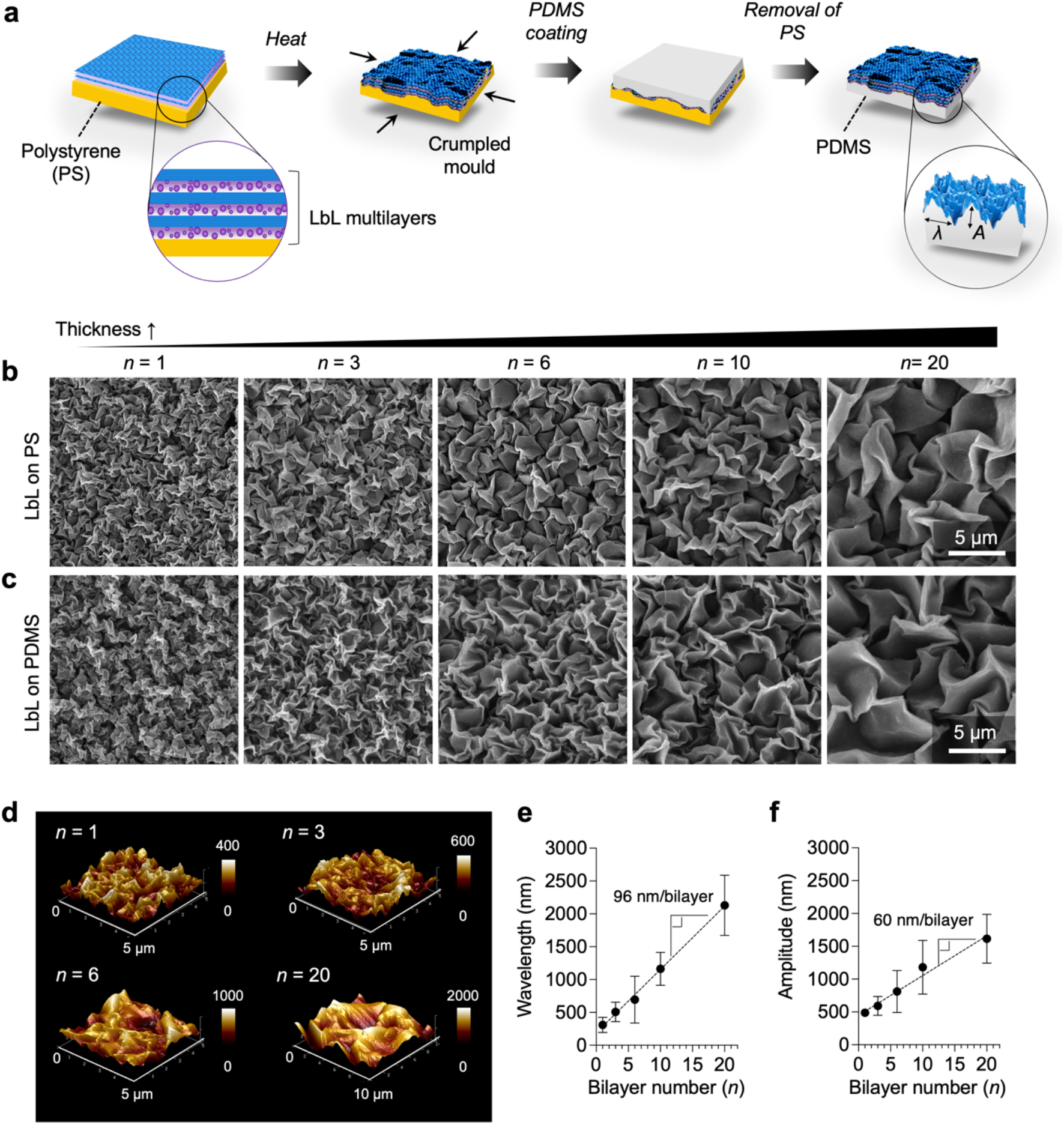
Morphological characterization of stretchable (MXene/GS)_*n*_ multilayers with crumple structures. **a**. Schematic of crumpled (MXene/GS)_*n*_ formation on PS films and PDMS as a stretchable substrate. The insert shows the increase in thickness by increasing the number of bilayers. SEM images of **b**. the crumpled (MXene/GS)_*n*_ multilayers on PS substrates and **c**. the crumpled multilayers transferred onto flexible PDMS substrates. **d**. 3D height AFM images of crumpled (MXene/GS)_*<u>n</u>*_ multilayers on PS substrates. **e**. Wavelength and **f**. peak-to-peak amplitude of the crumpled (MXene/GS)_*n*_ multilayers as a function of bilayer number (*n*).

Morphological details of the crumpled MXene LbL films were explored by SEM and AFM. SEM images show that sharp edges of the MXene wrinkles were generated and crumpled at different length scales and density. The crumpled MXene LbL film could be successfully transferred to a PDMS substrate. The structure comparison via SEM confirms the conformal transfer of the crumpled MXene LbL layers from the PS substrate to the PDMS substrate without obvious morphological distortion (**Fig. 2b-c**). The crumpled structures can be typically characterized with wavelength (*λ*, distance between peak to peak of periodic ripples) and amplitude (*A*, height difference between peak to peak)^[26]^ (**Fig. 2a**). These characteristic parameters can be tuned by controlling the thickness of the MXene multilayer coating at a high deformation strain (ε = 0.9). According to the classical buckling theory, a thin stiff coating with thickness *h* deposited on a uniaxially prestretched substrate will buckle with a characteristic wavelength of 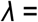 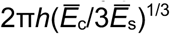 where the plane-strain elastic modulus 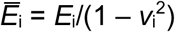 is given in terms of the Young’s modulus *E* and Poisson’s ratio *v*_i_ of the multilayer coating (c) or PS substrate (s), respectively^[27]^. This theory gives qualitative insight into the isotropic crumpled structures formed here through biaxial deformation. In this work, we analyzed SEM and AFM data to estimate the characteristic features of the crumpled MXene LbL films with different number of bilayers *n* (**Fig. 2d)**. For amplitude, we assumed the ridge-like structure of the MXene LbL film has a triangle waveform and used the relationship: 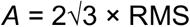 (root-mean-square). Our data show that as the film thickness increases from 4 nm (for *n* = 1) to 91 nm (for *n* = 20), the feature sizes increase nearly linearly: (i) *λ* from 308 nm to 2130 nm and (ii) *A* from 477 nm to 1615 nm (**Fig. 2e-f**). Fabrication of these different types of nanostructured MXene LbL films with finely crumpled configuration was scalable and robust.

To ensure long-term functionality of the engineered surfaces, we evaluated the mechanical stability of MXene multilayers on PS or PDMS substrates by characterizing interfacial failure under ultrasonication. The planar MXene multilayers on a PS substrate underwent interfacial delamination after ultrasonication for 15 min in water. In contrast, the crumpled MXene multilayers remained intact after ultrasonication for 15 min and showed negligible morphological distortion (**Fig. S4**). Furthermore, MXene multilayers on a PDMS substrate can withstand and remain intact without interfacial failure after 100 cycles of bending (radius of curvature ~2 mm) and stretching (~40% tensile strain) in water.

### Bactericidal effect of crumpled MXene coatings

**Figure 3a** illustrates the anticipated scenarios following bacteria inoculation on different surfaces. Substratum roughness, or the micro–nanotopography, is one of the most important surface characteristics for control of microbial attachment and the initial stage of biofilm formation^[28]^. Recent studies have shown that bacteria preferentially colonize recessed regions of the surface such as grooves with dimensions larger than themselves but fail to attach to surfaces with irregularly spaced pits with nanoscale feature sizes^[4c]^ (**Fig. 3a**). The effective dimensions of previously reported mechano-bactericidal nanofeature arrays are confined within a wavelength (spacing) range of 9–380 nm and an amplitude (height) of 100–900 nm^[4c]^. Based on these recent findings, we hypothesized that nano-crumpled structures of MXene multilayers with sufficiently dense spacing could impart the surfaces with bactericidal properties.

**Figure 3.**
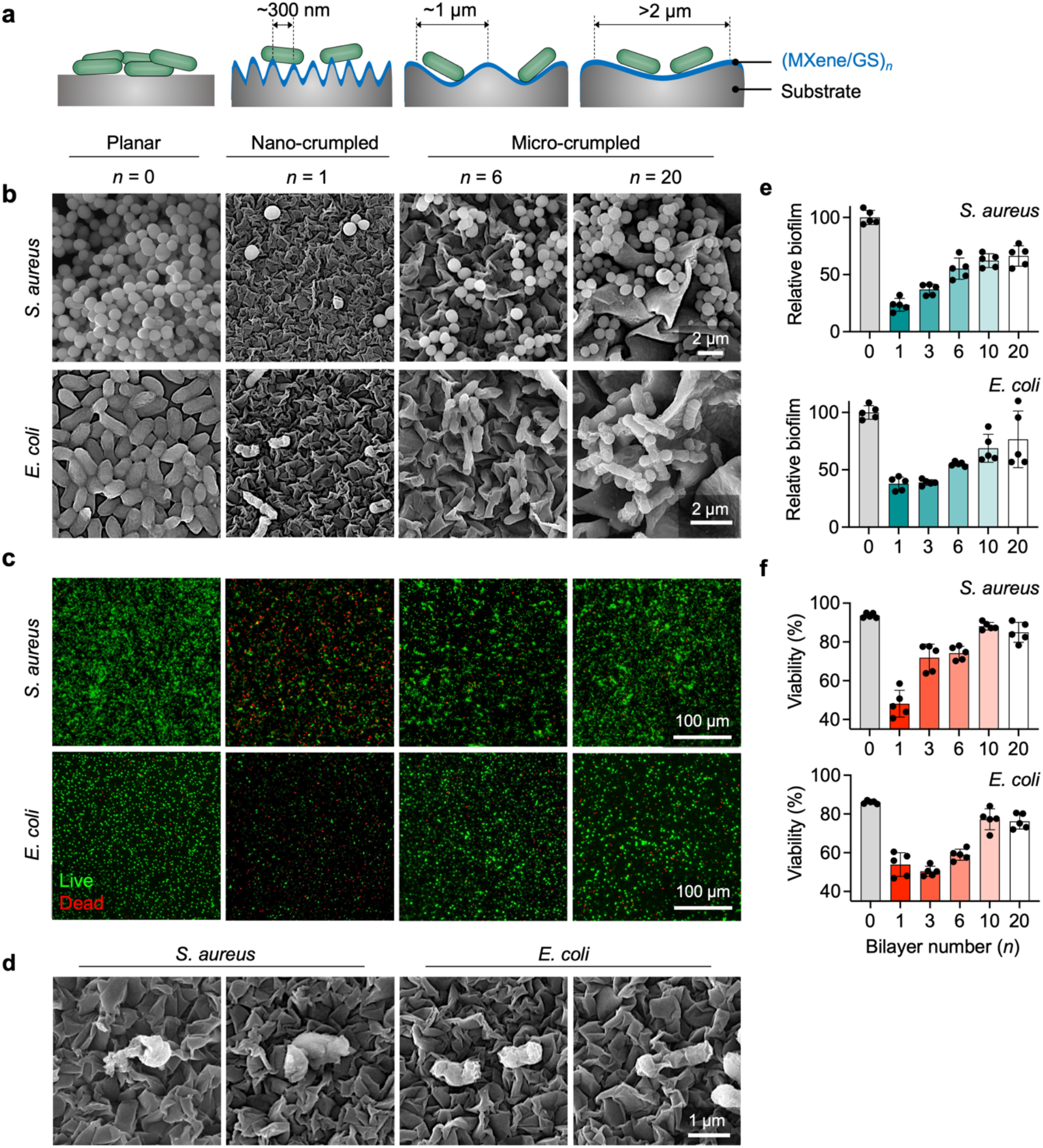
Comparison of antibacterial efficacy of crumpled (MXene/GS)_*n*_ multilayer films with different topographic parameters (wavelength, amplitude). **a**. Schematic illustration of the preferential attachment of bacteria to surface grooves of >2 μm and 1 μm to the tips of dense nanopillars of spacing 200 nm. **b**. SEM images of *S. aureus* and *E. coli* biofilm grown on flat controls and crumpled (MXene/GS)_n_ multilayers on PS substrates with different number of bilayers (*n*). **c**. Fluorescence images of *S. aureus* and *E. coli* biofilm grown on crumpled (MXene/GS)_n_ multilayers. Biofilm cells were labeled with STYO9 (green) and PI (red) for live and dead cells, respectively. **d**. SEM images of *S. aureus* and *E. coli* cells attached on crumpled (MXene/GS)_1_ surface. **e**. Relative biofilm coverage was calculated as total number of cells per unit area for *S. aureus* and *E. coli*, normalized to the flat controls. **f**. Cell viability was analyzed on the fluorescent images using Cellprofiler.

To test our hypothesis, we evaluated the antibacterial performance of the crumpled MXene multilayers with different *n*, using two bacterial species: (i) a Gram-positive *S. aureus USA300*, one of the most common causes of prosthetic valve endocarditis or catheter associated bloodstream infection^[29]^ and responsible for about one third of surgical-site infections^[30]^; and (ii) a Gram-negative *Escherichia coli UTI89*, the primary causative agent of catheter-associated urinary track infections (CAUTI)^[31]^. The antibacterial efficacy of the crumpled (MXene/GS)_*n*_ film against *S. aureus* and *E. coli* was evaluated by exploring the activity of the crumpled film directly using (i) SEM analysis (**Fig. 3b**) and (i) fluorescent microscopy with live/dead staining assay (**Fig. 3c**).

Overall, our results showed that the crumpled MXene surfaces have thickness-dependent (i) inhibition efficacy and (ii) bactericidal ability against cells that encounter the surface. SEM images showed that bacteria can form an intact and dense biofilm on a flat surface. These structures represent a mature biofilm structure in which the bacteria are closely interlinked and become highly resistant to immune killing and clearance and to treatment with antimicrobial agents^[32]^. On the other hand, bacteria with disrupted cell membranes could be found on the surface of the crumpled MXene multilayers with a lower number of *n* (**Fig. 3d**). The relative biofilm coverage, calculated as total cell count per unit area and then normalized to the flat control group, was found to decrease with the decrease in the thickness (or the number of bilayers) of the crumpled MXene multilayers. The (MXene/GS)_1_ multilayer with the smallest feature sizes (wavelength ~ 300 nm, amplitude ~ 500 nm) showed the best antibacterial activity with the relative biofilm coverage of 23 ± 5.6% and 37 ± 5.8% (**Fig. 3e**) for *S. aureus* and *E. coli*, respectively. This result is congruent with the effective dimensions of previously reported mechano-bactericidal nanofeature arrays^[4c]^. For the groups with larger *n*, we observed that bacteria could colonize the recessed regions of the crumpled surface with dimensions larger than themselves (*i.e*., wavelength > 1 μm). Fluorescence images of biofilms labeled with STYO9 (live stain) and propidium iodide (PI; dead stain) showed that the flat surface was covered by live bacteria with a dense surface coverage, while many dead bacteria with some live cells with a low surface coverage could be found in the crumpled (MXene/GS)_1_ group. The bacteria cell viability on the (MXene/GS)_1_ multilayer was found to be 48 ± 6.9% and 54 ± 6.1% for *S. aureus* and *E. coli*, respectively (**Fig. 3f** and **Fig. S5**). The cell viability increases from *n* = 6 to 10 for *E. coli* (58 ± 2.8% vs. 77 ± 5.4%, *p* < 0.0001) and from *n* = 1 to 3 for *S. aureus* (48 ± 6.9% vs. 72 ± 6.9%, *p* < 0.0001). These transitions occur when the characteristic wavelength of the crumple structures becomes compatible to the size of bacteria (*S. aureus*, 640 ± 62 nm and *E. coli*, 1,500 ± 290 nm in long axis, **Fig. S6**), which is congruent with the recent finding^[4c]^. Taking into consideration both biofilm coverage and cell viability, we found that the crumpled Mxene multilayers could reduce biofilm biomass up to 89% and 81% for *S. aureus* and *E. coli*, respectively, compared to the flat control. To test release of film components, we performed the Kirby-Bauer disk diffusion assay using crumpled MXene LbL films that provides qualitative information regarding the amount of GS or MXene that has diffused through agar by measuring the clear zone of inhibition (ZOI). We observed no ZOI even in the (MXene/GS)_20_ group against *S. aureus*, suggesting no or negligible release of the film components (**Fig. S7**)

### Dynamic deformation by external stimuli

To examine the effect of surface deformation on biofilm detachment, we conducted experiments with biofilm formation of *S. aureus* and *E. coli* on the crumpled (MXene/GS)_1_ multilayers on PDMS substrates. We hypothesized that a small surface deformation by mechanical stretching would exert enough force to detach the attached cells including the debris of dead bacterial cells. We first cultured bacterial cells on different surfaces for 24-48 h without perturbation, and then we applied on-demand stretching (uniaxial 20% strain for 30 cycles in 3 min) only to the dynamic group. After on-demand stretching, we performed live/dead fluorescence microscopy. As expected, our results showed that the dynamic deformation to a prescribed strain (20%) disrupted biofilms of both species (**Fig. 4**). We first validated that the crumpled (MXene/GS)_1_ on PDMS substrates exhibit similar antibacterial efficacy as the original films on PS substrates. Specifically, the relative biofilm coverage deceased from the flat PDMS (flat control) to the crumpled MXene film (static control): (i) 100 ± 17% vs. 30 ± 8.4% (*p* < 0.0001) for *S. aureus* and (ii) 100 ± 18% vs. 40 ± 8.0% (*p* < 0.0001) for *E. coli*. Strong effects were observed for the removal of adhered bacterial cells (cultured for 24-48 h without actuation first) by activating mechanical stretching on-demand at 20% strain for 3 min. This treatment caused biomass reduction of 97% for *S. aureus* and 99% for *E. coli*, compared to the static control: (i) 30 ± 8.4% vs. 1.1 ± 1.0% (*p* = 0.0016) for *S. aureus* and (ii) 40 ± 8.0% vs. 0.36 ± 0.35% (*p* = 0.0013) for *E. coli*. Compared to the flat control, the crumpled MXene multilayers with dynamic topography via on-demand stretching could reduce biofilm biomass nearly completely (≥99%) for both species.

**Figure 4.**
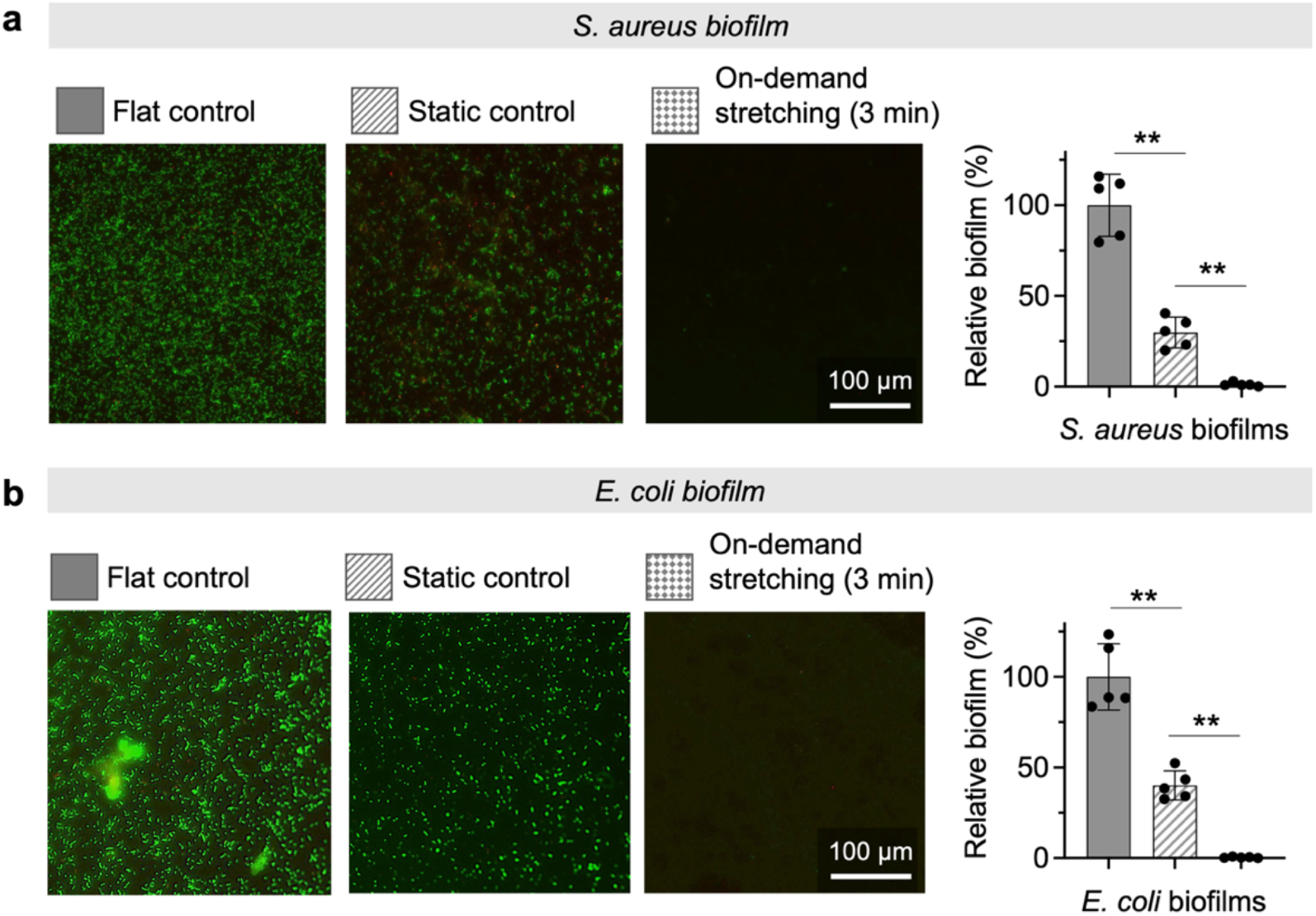
Fluorescence images of biofilm grown on (i) flat controls, and (ii-iii) crumpled MXene multilayers (MXene/GS)_1_ with or without dynamic deformation after incubating for 24 h and 48 h for **a**. *S. aureus* and **b**. *E. coli*, respectively. After overnight bacterial growth, on-demand dynamic topographies were operated by applying 20% strain for 30 cycles in 3 min. Relative biofilm coverage was calculated as total number of cells per unit area for *S. aureus* and *E. coli*, normalized to the flat controls.

### Safety to mammalian cells

To evaluate the clinical potential of this design, we further tested the biocompatibility of the crumpled MXene films to mammalian cells. We performed the cell viability test on individual components and confirmed that there is no apparent cytotoxicity associated with the individual component at the concentrations used in this study (**Fig. 5**). We focused on investigating the effects of nano- and micro-structured 3D MXene surfaces on cell growth of mammalian cells. We cultured murine fibroblast NIH-3T3 cells on the crumpled MXene LbL films with varying number of bilayers (*n* = 1 to 20). The results showed that the nano-/micro-structures of crumpled LbL films do not affect cell viability. A sharp transition in the number of adhered cells on films was observed from *n* = 10 to *n* = 20.

**Figure 5.**
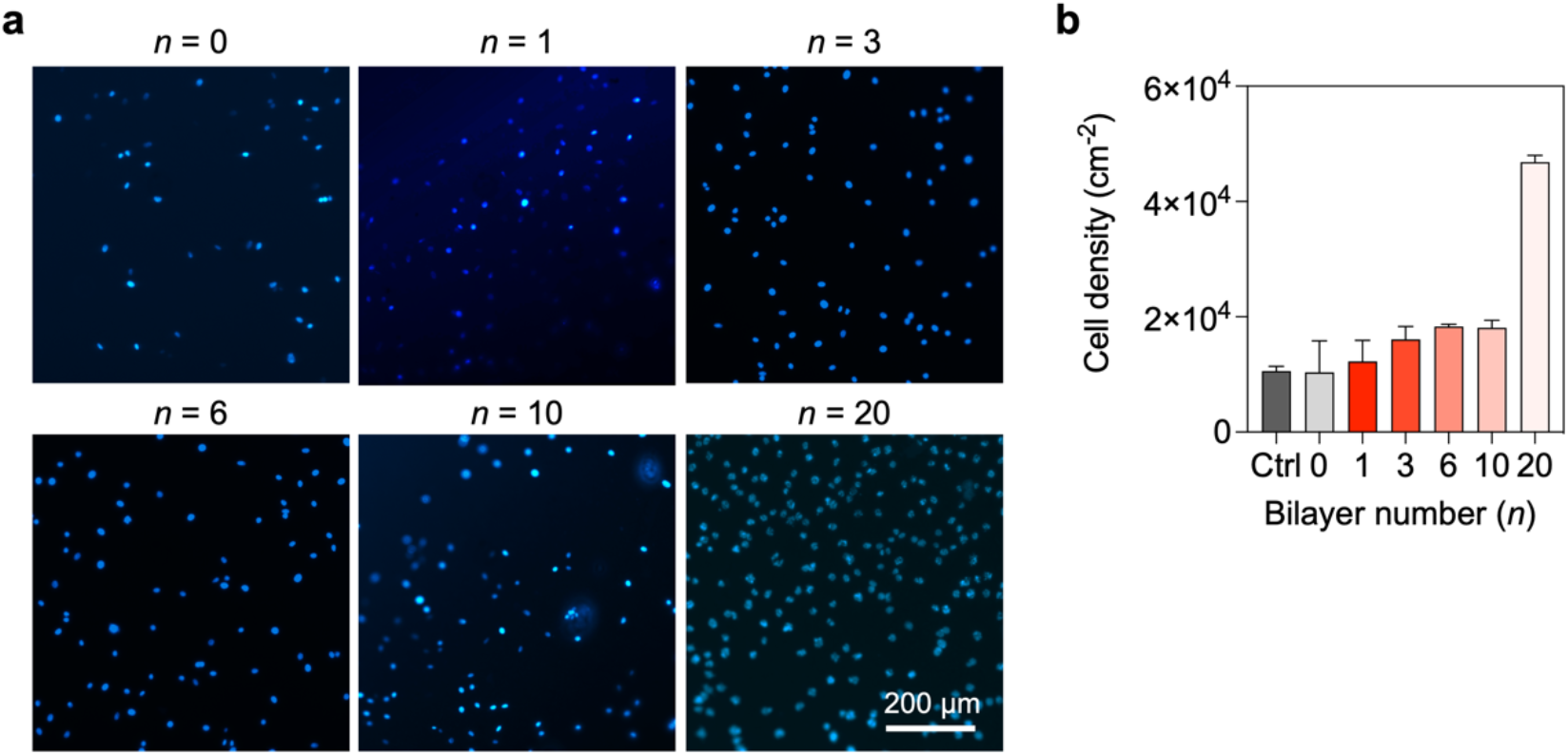
Biocompatibility testing of crumpled structures. **(a)** Effects of crumpled structure on mammalian cells (fibroblast NIH/3T3) grown on different (MXene/GS)_*n*_ multilayers for 24 hours. **(b)** Cell density was calculated as the number of cells per unit area (cm^2^) to compare the growth of mammalian cells on the plane well-plate (Ctrl), the flat control (*n* = 0) and (MXene/GS)_*n*_ multilayers.

## DISCUSSION

Antibiotic resistance is a global human health threat, causing routine treatments of bacterial infections to become increasingly difficult. The problem is exacerbated by biofilm formation by pathogenic bacteria on the surfaces of indwelling medical devices that facilitate high levels of resistance to antibiotics. Recently, the development of new nature-inspired nano/micro-structured surfaces has been gaining popularity due to their mechano-bactericidal or antibiofouling activity. A potential drawback to static topographies is that effective biofilm control depends on the direct interaction between bacteria and the surface. Although static topographies with specific micron or nano-scale features may initially prevent bacterial adhesion and biofilm formation, cells that manage to attach tend to multiply and overcome these features over time. For surfaces that have bactericidal effects, it is also possible that dead cell debris may protect other cells that attach to them rather than the original topographic features.

To address these limitations of static topographies, some platform technologies based on dynamic topographies driven by mechanical stretching^[33]^, pneumatic actuation^[34]^, electrical voltage^[33a]^, or magnetic actuation^[35]^ have been developed for sustained antifouling activities. These recent studies have revealed that (1) surface deformation by the applied strain exceeding threshold values (5-15%) leads to significant detachment of bacterial biofilms, and (2) on-demand actuation of surface deformation is sufficient for removal of established biofilms. Although these recent studies marked a step forward and opened a new field of dynamic topography, these designs lack bactericidal functionality, thereby necessitating a subsequent delivery of bactericidal agents (antibiotics) to sensitize the detached bacteria cells. For optimal long-term antibacterial performance in biomedical implants, there is a need for multifunctional platform technologies that: (1) inhibit bacterial attachment, (2) mechanically kill cells that encounter the surface, and (3) detach dead cell debris to restore functionality.

Here we demonstrated a new efficient strategy for fabricating stretchable MXene multilayers with finely tuned crumple structures for long-term biofilm control and explored the interaction of the structured surfaces with two different species of bacteria (*S. aureus* and *E. coli*). The stretchable MXene multilayers with crumpled structures have many noteworthy features: (1) They show a linear increase in thickness with the bilayer number with a high resolution of 4.3 nm/bilayer, indicating that our LbL self-assembly method is very precise. (2) The feature sizes of crumpled surfaces also increase linearly with the thickness of as-prepared multilayers, and thus they can be finely tuned by simply controlling the number of layers. (3) The crumpled MXene multilayers show excellent mechanical durability. (4) Nanoscale geometry of MXene multilayers significantly affects their antimicrobial activity. Our results demonstrated that the static MXene multilayers have thickness-dependent antibacterial efficacy against bacteria. Based on the results, we can explain the relationship between antibacterial activity and surface feature sizes of the crumpled MXene multilayers as follows. In addition to the mode of antibacterial activity mediated by direct contact to MXene^[16, 36]^, our approach reveals a potent antibacterial effect of the geometrically tuned MXene surfaces that do not release any bactericidal substances. The finely tuned surface crumples were composed of peaks and valleys, and the sharp edge of the peaks can inhibit attachment of bacteria cells and the abundance of sharp edges with better charge transfer in the MXene nanosheets can cause cell membrane destruction^[37]^. (5) Dynamic deformation of MXene multilayer surfaces could induce significant detachment of bacterial biofilms (>99%) by applying 20% strain (30 cycles for 3 min) and restore clean surfaces with multifunctionality. (6) Crumpled structures are safe towards mammalian cells with no apparent cytotoxicity, physical damage, or reduction in cell viability. Interestingly, we observed a sharp transition in the number of adhered cells on films from *n* = 10 to *n* = 20. This increase in the number of cells by a factor of four is likely due to the increase in the effective surface area for cells to grow. (7) They have surface-agnostic features and excellent scalability. Overall, our design meets the conditions required for biomedical implants, namely cytocompatibility, durability, proven bactericidal and antifouling effects, practicality, low-cost and scalability^[4c]^.

The current study had some limitations. First, the current design is not readily applicable to fouling control *in vivo* due to limitations of the actuation mechanism, which is mechanical stretching. Further investigation into alternate substrates, such as thermo-responsive polymers or elastomers, that can expand and contract upon a change in applied temperature^[33b]^ or pressure^[34b]^ may be more amenable to *in vivo* applications. Second, there is room for improved antibacterial activity of the static MXene multilayers. Our study was designed primarily to explore the contact-based physical interaction between bacteria and geometrically tuned MXene surfaces. We can improve the antibacterial efficacy by further modifications such as (i) conjugating metal oxides to MXene nanosheets, which can induce oxidative stress, or (ii) applying NIR light (808 nm) for photothermal therapy. MXene has an excellent light-to-heat conversion efficiency (~100%)^[38]^, and thus MXene multilayers can be used for synergistic mechano-bactericidal and photothermal killing as a potent antibacterial strategy. These additional engineering aspects are straightforward to implement without altering the system design and fabrication method.

In summary, we developed a new antibacterial strategy based on the geometrically tuned MXene surfaces with dynamic topographies that do not release bactericidal substances. We believe that our approach validates a promising 2D material MXene-based multilayers as an efficient platform with long-term antibacterial properties that can be fabricated in a low-cost, scalable manner and be applicable for a range of biomaterials and implantable devices. Further studies using animal models will help validate the clinical potential of this technology, which is part of our future work.

## METHODS AND EXPERIMENTAL

### Materials

Gentamicin sulfate (GS), anhydrous ethanol, acetone, dichloromethane (DCM), (tridecafluoro-1,1,2,2-tetrahydrooctyl)trichlorosilane, lithium fluoride (LiF), sodium alginate, hydrochloric acid (HCl), sulfuric acid (H_2_SO_4_) were purchased from Sigma-Aldrich. PDMS prepolymer and curing agent (Sylgard 184) were purchased from Fisher Scientific. Ti_3_AlC_2_ MAX phase powders were purchased from Laizhou Kai Kai Ceramic Materials Co. (300 mesh, ≥99%). Clear heat shrinkable polystyrene (PS) films were purchased from Grafix. All water was deionized (DI) (18.2 megohm, Mill-Q pore). All reagents were used as received without further purification.

### Synthesis of Ti_3_C_2_T_x_ MXene

Ti_3_C_2_T_x_ MXene nanosheets were prepared by in situ HF etching according to the previous literature with some modification^[19]^. Max phase Ti_3_AlC_2_ powder (1.0 g) was etched with a solution of LiF (7.5 g) and HCl (9M, 40 mL) under stirring at 35°C for 24 h to remove its Al layers. The solid residue was obtained by centrifugation and washed with DI water several times until the pH value reached around 6.0. Subsequently, 100 mL of DI water was added to the residue, and the mixture was bath sonicated for 1 h at 4°C and centrifuged at 3500 rpm for 30 min to separate the heavier components. The supernatant contained the stable Ti_3_C_2_T_x_ MXene nanosheets. The concentration of obtained MXene suspension was ~10 mg mL^−1^.

### LbL assembly of MXene multilayers

Multilayers were deposited at the surface of various substrates: PS films, slide glass, silicon wafer, and PDMS. The GS and MXene sheets were diluted to a concentration of 1.0 mg/ml in DI water. The pH values of the GS solution and MXene dispersion were 5.0 and 6.1, respectively, and both solutions were used without adjusting the pH. All substrates were cleaned by sequential sonication in ethanol and water for 15 min each. After washing, the glass was dried with nitrogen and treated with oxygen plasma (Tergeo-Plus, PIE Scientific) for 10 min at 50 W under vacuum. For the dip-based multilayer assembly, we used an automated dipping robot (KSV NIMA Dip coater, Biolin Scientific) and the Ti_3_C_2_T_x_ MXene and GS solutions with a concentration of 1 mg mL^−1^.

The plasma-treated substrates were first immersed in PDAC solution for 30 min to deposit a base layer, and rinsed with DI water twice for 2 min each to remove the weakly bound molecules. After that, bilayer multilayers were fabricated by alternate dipping in a solution of cationic species for 5 min followed by two consecutive rinse steps in DI water baths twice for 2 min each, and then into anionic species for 5 min followed by the same rinse cycle. This cycle made one bilayer of (MXene/GS)_1_, and the bilayer was repeated to fabricate the desired multilayers, denoted as (MXene/GS)_*n*_ where *n* is the bilayer number. The as-prepared multilayer films were dried at room temperature and stored in a closed container prior to subsequent analysis.

### Fabrication of crumpled MXene multilayers on PDMS substrates

As prepared MXene multilayers on PS substrates were cut into a desired size and shrunk by heating in an oven at 130 °C for 5 min to form crumpled films. The strain of the PS film was determined using the equation ε = (*L*_i_ – *L*_f_)/*L*_i_. Subsequently, a solution of degassed PDMS prepolymer and curing agent (10:1, w/w) was spin-coated (500 rpm, 30 s) onto the crumpled MXene multilayers (thickness of ~400 μm) and then cured at 60 °C for 4 h to generate PS/MXene LbL/PDMS films. The PS/MXene LbL/PDMS films were immersed in dimethylformamide (DMF) for 1 h to remove PS and then washed in acetone and ethanol. The MXene multilayers on the PDMS substrate were obtained after drying at room temperature.

### Material characterization

Film thickness and surface roughness were determined by AFM (Dimension Icon, Bruker Corp.) in tapping mode. Film deposition was measured using a UV-Vis spectrophotometer (Agilent 8453). MXene multilayers grown on silicon wafers were scratched with a razor blade, and thickness was measured at three predetermined locations. The surface morphology of the LbL films were observed using SEM (Tescan Mira3). The zeta potential of MXene dispersions at pH 6 was measured by a Zetasizer (ZEN3600, Malvern Instruments Ltd.). XRD spectra were obtained from the MXene multilayers assembled on glass slides and were conducted on an X-Ray diffractometer (Rigaku Ultima IV) in the range of 5–30°. The interlayer spacing *d* (nm) of MXene multilayers was calculated according to the Bragg’s law^[39]^:

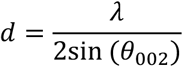

where *λ* (*λ* = 0.15406 nm) is the wavelength of X-ray used, and *θ*_002_ is the scattering angles of the (002) peak of MXene multilayers.

### Antibacterial assay

*S. aureus* USA300 and *E.coli* UTI89 were used as the model microorganisms in this study. To test the performance of surface topographies in antibacterial activity, crumpled or planar surfaces (1 × 1 cm^2^) were attached on the bottom of a 24-well plate. Bacterial cells were first grown in Tryptic Soy medium (for *S. aureus*) or LB medium (for *E.coli*) overnight at 37 °C with shaking at 200 rpm. Overnight cultures were diluted to ~10^4^ CFU mL^−1^ and inoculated on the surfaces for biofilm growth. The surfaces were incubated at 37 °C for 24 and 48 h for *S. aureus* and *E. coli*, respectively.

To quantify the attached cells, we labeled the cells with a mixture of SYTO9 and propidium iodide (PI). Before staining, the biofilm surfaces were washed three times with DI water and then labeled with a mixture of SYTO9 (5 μM) and PI (20 μM) for 15 min at room temperature in the dark. The surfaces are imaged using an Axio Observer fluorescence microscope (Carl Zeiss Inc.,). Fluorescence images were taken from randomly selected positions (*n* ≥ 3) on each surface. Image analyses were performed using ImageJ and CellProfiler. The imaging channels were used to count live and dead cells.

To test the removal of adhered bacterial cells via dynamic topography, biofilms were formed on crumpled surfaces at 37 °C with an inoculation concentration of ~10^4^ CFU mL^−1^. Before on-demand actuation, the biofilm surfaces were washed and then stretched using the method as previously described^[33a]^. Specifically, the multilayer coated PDMS substrate with biofilm was carefully transferred to a caliper, was stretched uniaxially to a prescribed strain (~20%) using a caliper for 30 cycles within 3 min in DI water (**Fig. S8**). After the on-demand biofilm removal, the surfaces were washed three times with DI. Then the remaining bacterial cells were labeled and imaged as described above. The biofilm with and without the dynamic actuation was quantified in the same way as described above for on-demand actuation.

### Biofilm imaging using SEM

Biofilms on MXene surfaces with static crumples were formed as described above. Biofilm samples were fixed in 2.5% glutaraldehyde in cacodylate buffer (CB) overnight at 4°C. The samples were then incubated in 1% osmium tetroxide-CB for secondary fixing and staining for 1 h. Thorough washing was done between steps. The fixed samples were dehydrated using 10, 30, 50, 70, 80, 90, 95, and 100% ethanol for 15 min each. The 100% ethanol step was repeated three times. The dehydrated samples were dried using a critical point dryer (Leica EM CPD300). The samples were sputter coated with gold, and then imaged using the Tescan Mira3 SEM.

### Biocompatibility testing

NIH/3T3 mouse fibroblasts cells were used to investigate the biocompatibility of the crumpled topographies. Briefly, NIH/3T3 cells were cultured in growth medium (DMEM supplemented with 10% FBS and 1% of antibiotic and antimycotic solution) in a humidified incubator (37 C; 5% CO2 in air). Cells were subcultured when near 80% confluence with the use of 0.05% Trypsin-EDTA. All cells used in these studies were passage number 5-15.

After overnight exposure (18-24 h) to different concentrations of the individual components or crumpled surfaces, cells were examined by using the CellTiter-GlLuminescent Cell Viability Assay (Promega, Madison, WI) and the conventional DAPI staining. CellTiter-Glo luminescent assay is a method to determine the cell viability based on quantitation of the ATP present, which signals the presence of metabolically active cells, and DAPI staining is for cell counting by fluorescent imaging. The viability assays were performed according to the manufacturer’s specifications.

### Statistics

Statistical analyses and data plotting were performed in GraphPad Prism 9. Data are reported as mean +/− standard deviation of a minimum of 3 samples. For correlations, the linear least squares fitting was performed at the 95% confidence level, and the Pearson correlation coefficient (*r*) was used to quantify the correlations between different variables. Group differences were tested using the nonparametric Mann-Whitney test for two groups and analysis of variance (ANOVA) with post hoc analysis for more than two groups. All tests were two-sided, and *p* < 0.05 was considered statistically significant.

## Data availability

The data that support the findings of this study are available from the corresponding author upon reasonable request.

## Supplementary Materials

Fig. S1: MXene synthesis and characterization.

Fig. S2. Cytotoxicity of individual components.

Fig. S3. Self-assembly formation of MXene/GS hybrids.

Fig. S4. Mechanical stability testing.

Fig. S5. Biofilm measurements via Live/Dead assay.

Fig. S6. Size distribution of bacteria cells

Fig. S7. Comparison of antibacterial activity via Kirby-Bauer diffusion assay

Fig. S8. Experimental setup for dynamic testing.

## Acknowledgements

This work was supported by a Startup Grant from University of Michigan, College of Engineering. We would like to thank Zhongrui (Jerry) Li for assistance with XRD measurements, and Haiping Sun for assistance with SEM. The authors greatly appreciate the use of equipment available in BioInterface Institutes for confocal imaging.

## Author contribution

J.M., conceived the study. N.N., M.A., J.M. performed experiments. N.N., M.A., P.Y.C., J.S.V., J.M. analyzed data. N.N., M.A., P.Y.C., J.S.V., J.M. wrote the manuscript, which was edited by all authors.

