## Supplementary materials for "Stretchable, nano-crumpled MXene multilayers impart long-term antibacterial surface properties"

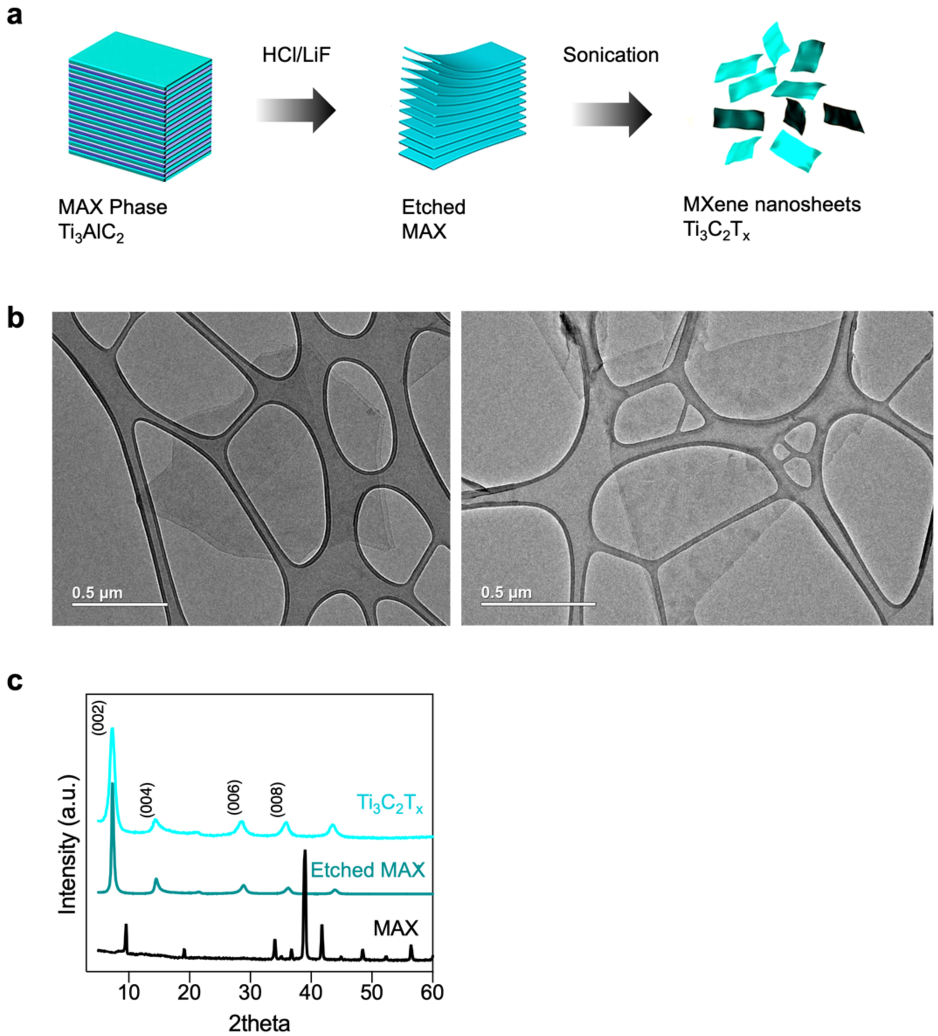


**Figure S1. MXene synthesis and characterization.** (a) Schematic illustration of $Ti_3C_2T_x$ MXene synthesis. Selective etching of Al atoms from $Ti_3AlC_2$ MAX phase in a LiF and HCl mixture. (b) TEM images of $Ti_3C_2T_x$ MXene nanosheets (c) XRD patterns of powder $Ti_3AlC_2$ (MAX phase), MXene before and after delamination, denoted as $Ti_3C_2$ and $Ti_3C_2T_x$, respectively. Multilayer and Nanosheet papers were produced by vacuum filtration of MXene solution. After exfoliation and delamination, the characteristic peak (002) of the $Ti_3AlC_2$ shifts from 9.5º to 6.8º for the MXene particles.

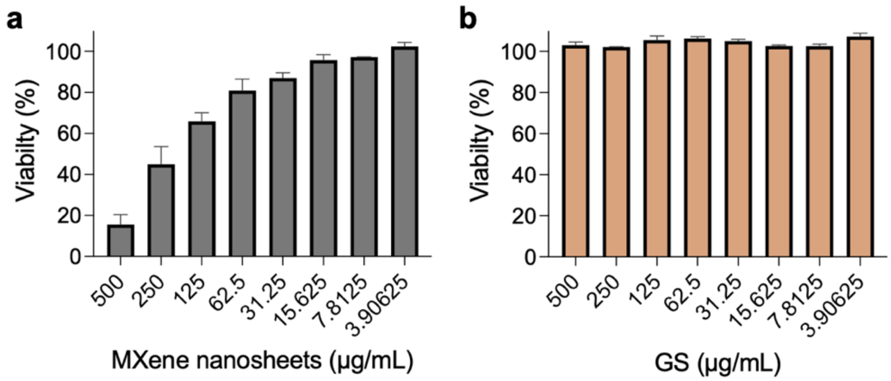


**Figure S2. Cytotoxicity of individual components**. Metabolic activity of mammalian cells (NIH/3T3) after 16–18 h treatment to MXene (**a**) or GS (**b**) at different concentrations. The luminescent values are normalized to negative control (*i.e.,* standard growth medium). MXene nanosheet dispersion with a concentration greater than 250 µg/mL are cytotoxic.

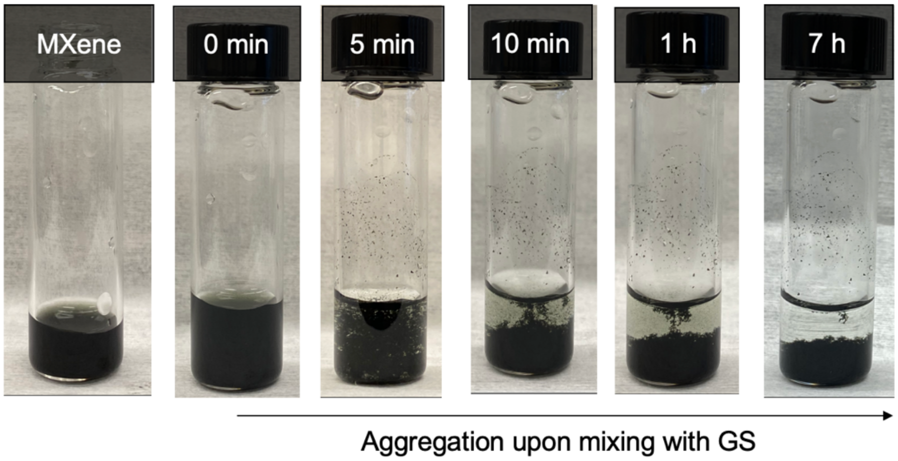


**Figure S3. Self-assembly formation of MXene/GS hybrids** after mixing the MXene dispersion with GS solution. The photographs are for 1 mg/mL MXene dispersion, and the MXene/GS hybrids (with 1 mg/mL GS) at the different time after mixing, from left to right.

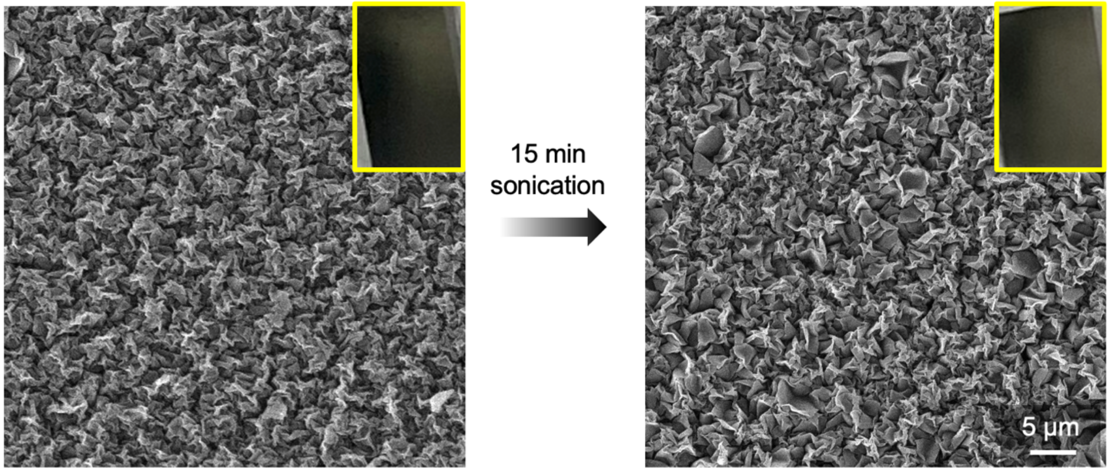


**Figure S4. Mechanical stability testing.** Representative SEM images of a crumpled (MXene/GS)$_6$ multilayer before and after 15 min bath sonication. No obvious morphological distortion was observed.

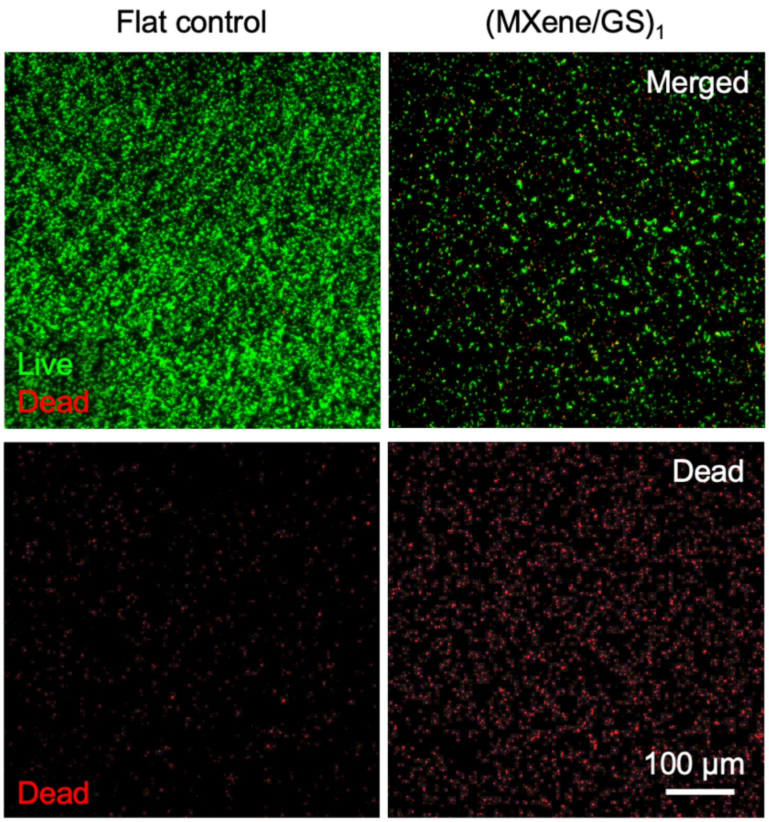


**Figure S5. Biofilm measurements via Live/Dead assay.** Merged (live and dead) and dead channels of fluorescent microscopic images of *S. aureus* biofilm grown on the flat control and the crumpled $(MXene/GS)_1$.

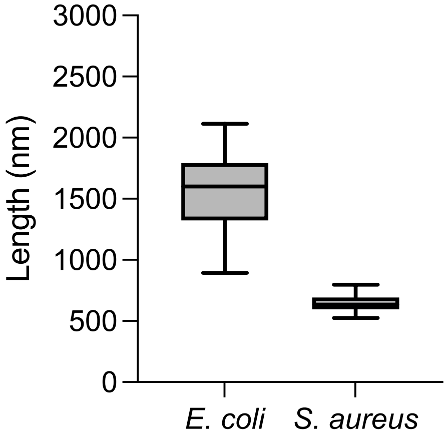


**Figure S6.** Size distribution of *S. aureus* and *E. coli* cells. Size of the long-axis was analyzed using SEM data.

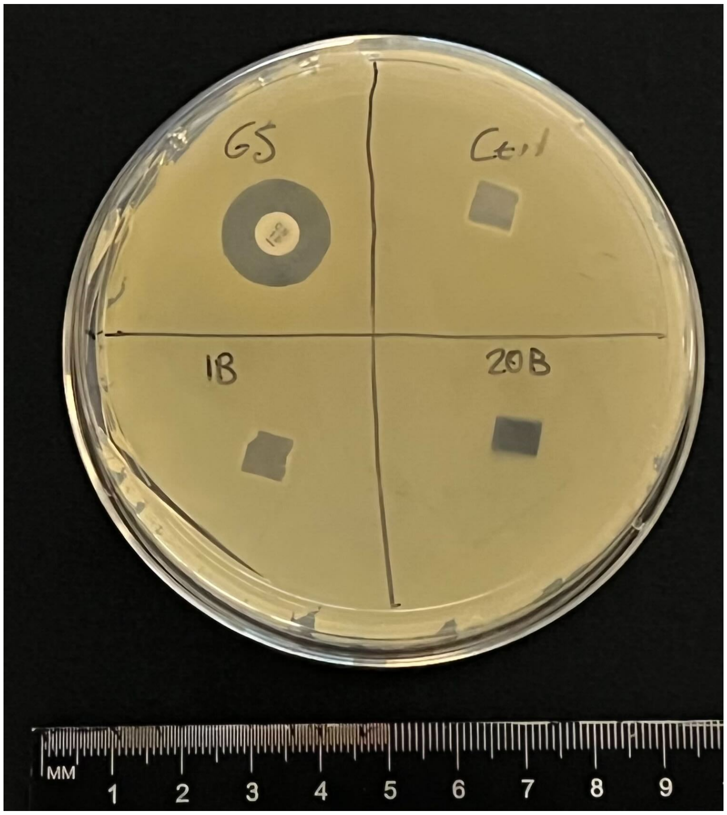


**Figure S7.** Comparison of antibacterial activity between various types of surfaces against *S. Aureus* via Kirby-Bauer diffusion assay. GS, commercially available BD Sensi-Disk control; Ctrl, uncoated substrate control; 1B, crumpled $(MXene/GS)_1$; 20B, crumpled $(MXene/GS)_{20}$ multilayers. This data confirms no or negligible release of the film components from the $(MXene/GS)_n$ groups.

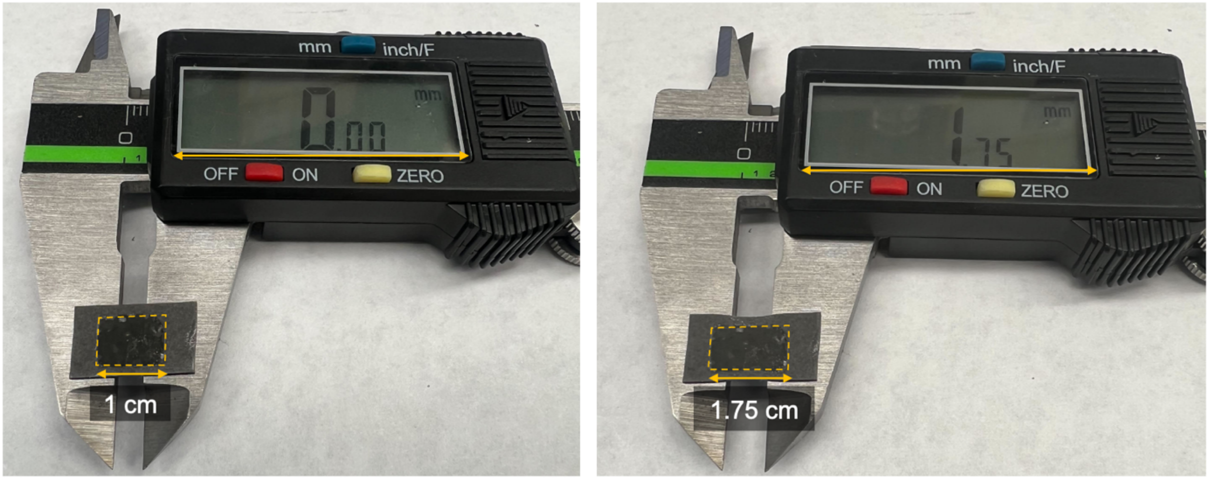


**Figure S8. Experimental setup for dynamic testing.** Digital photographs of the crumpled $(MXene/GS)_n$ multilayers on PDMS substrates, mounted on 3M VHB double-sided tape, in its original (left) and stretched (right) state. A caliper was used to control the applied strain to ~20%.
